# A 3500-year-old leaf from a Pharaonic tomb reveals that New Kingdom Egyptians were cultivating domesticated watermelon

**DOI:** 10.1101/642785

**Authors:** Susanne S. Renner, Oscar A. Pérez-Escobar, Martina V. Silber, Mark Nesbitt, Michaela Preick, Michael Hofreiter, Guillaume Chomicki

## Abstract

Domestication of the watermelon (*Citrullus lanatus*) has alternatively been placed in South Africa, the Nile valley, or more recently West Africa, with the oldest archeological evidence coming from Libya and Egypt. The geographic origin and domestication of watermelons has therefore remained unclear. Using extensive nuclear and plastid genomic data from a 3,560-year-old *Citrullus* leaf from a mummy’s sarcophagus and skimmed genomes for representatives of the seven extant species of *Citrullus*, we show that modern cultivars and the ancient plant uniquely share mutations in a lycopene metabolism gene (LYCB) affecting pulp color and a stop codon in a transcription factor regulating bitter cucurbitacin compounds. This implies that the plant we sequenced had red-fleshed and sweet fruits and that New Kingdom Egyptians were cultivating domesticated watermelons. The genomic data also identify extant Sudanese watermelons with white, sweet pulp as the closest relatives of domesticated watermelons.

**Significance statement:** With some 197.8 million tons in 2017, watermelon, *Citrullus lanatus*, is among the World’s most important crops, yet its area of origin and domestication have remained unclear, with competing hypotheses favoring South Africa, West Africa, Central Africa, or the Nile valley. We generated extensive nuclear and plastid genomic data from a 3500-year-old leaf from a Pharaonic sarcophagus and performed genome skimming for representatives of all other *Citrullus* species to compare key genes involved in fruit bitterness and color. White-fleshed, non-bitter melons from southern Sudan are the closest relatives of domesticated watermelon, and the ancient genome shares unique alleles with a red-fleshed, non-bitter domesticated form (but no wild forms), implying that 18^th^ Dynasty Egyptians were cultivating domesticated watermelon by 3500 years ago.

## Introduction

Among the World’s most important cucurbit fruit crops is the watermelon (*Citrullus lanatus*). With a production of 197.8 million tons in 2017, watermelon accounts for over a third of the global tropical fruit production (*1*). Its geographic origin and time of domestication, however, remain contentious (*2-7, supplementary text).* Competing hypotheses have favored South Africa (*7, 8*), Central Africa (*2*) or the Nile valley (*9-15*). More recently, molecular data have pinpointed West Africa (*16, 17*) as the region of the crop’s origin, yet archaeological evidence is restricted to Egypt and Libya. Besides *C. lanatus*, the genus *Citrullus* contains six other species, of which four (*C. amarus, C. ecirrhosus, C. naudinianus*, and *C. rehmii*) occur mostly in the Namib-Kalahari region, one (*C. mucosospermus*) in West Africa (Benin, Ghana, and Nigeria), and one (*C. colocynthis*) from northern Africa to west India (**Fig. 1A**) and naturalized also in Australia (*6, 17*). All have white pulp that cannot be eaten raw due to the presence of terpene bitter compounds called cucurbitacins. Only fruits of *C. mucosospermus* are sometimes not bitter, but instead bland-tasting (*18*). *Citrullus lanatus* itself is cultivated worldwide, but wild progenitors remained elusive despite the resequencing of 20 accessions from germplasm collections (*16*). Putative wild forms with small fruits and sweet white pulp have been reported since the 19^th^ century from the Kordofan region, a former province of Sudan, bordering North and South Darfur (*6, 9-15*) (**Fig. 1A, B**), but their genetic affinity to domesticated and other watermelons has not been tested. Moreover, no wild species of *Citrullus* with sweet pulp have been found in South Africa.

**Fig. 1.**
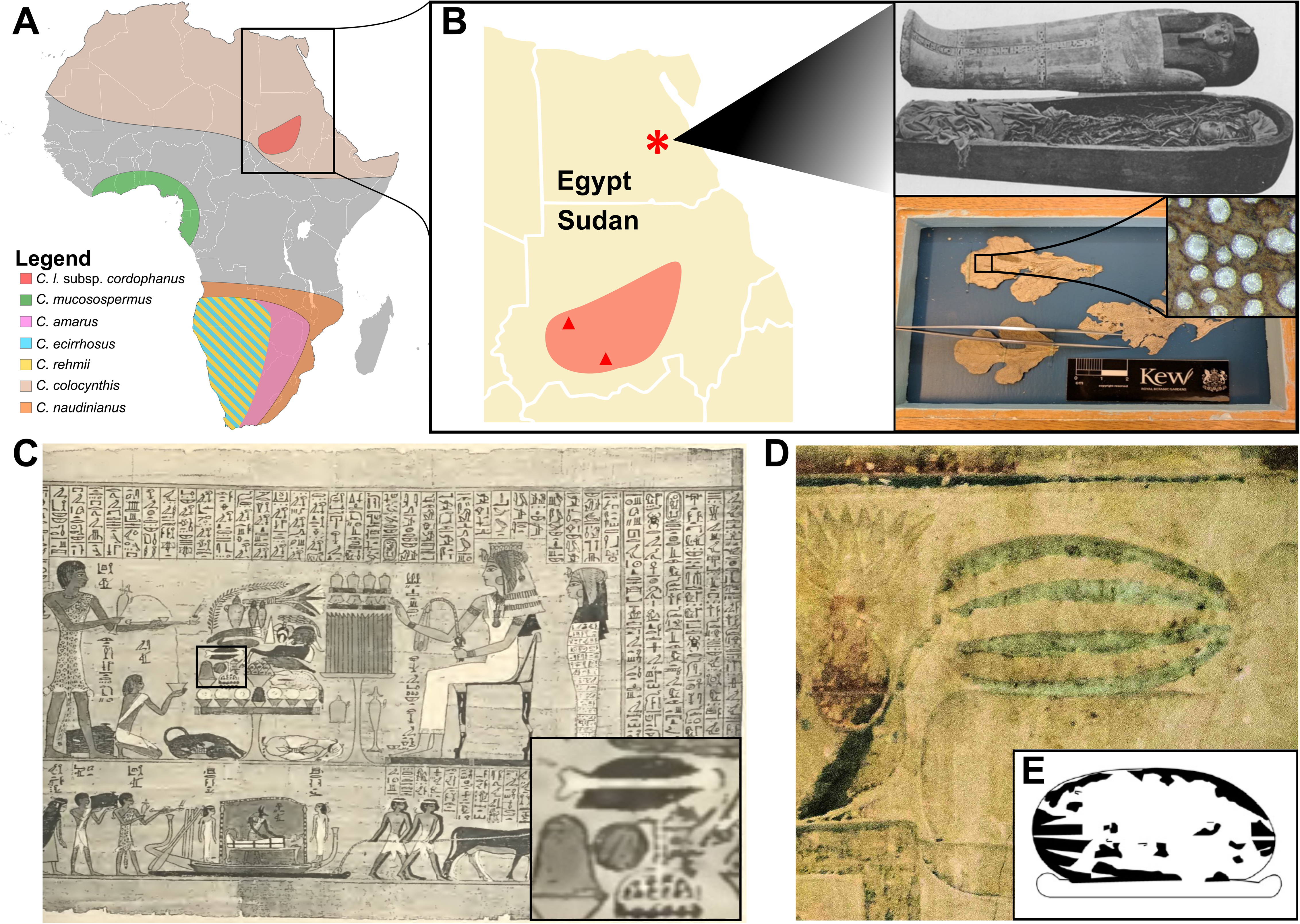
Distribution of *Citrullus* species, archaeological samples and illustrations of watermelon in Ancient Egypt. **(A)**, Map showing the distribution of *Citrullus* species in Africa (*C. colocynthis* extends to West India). (**B)**, An asterisk marks the location of the tomb at Deir el-Bahari near Luxor, Egypt, in which a leaf fragment was found on “Unknown Man C” (Cairo museum mummy CG61067) in the 19^th^ century and deposited in the Economic Plant Collection of Kew (***9-12***). Red triangles and patch mark the Kordofan region and locations with wild watermelons in North and South Darfur. Top, sarcophagus of Amenhotpou from Deir el-Bahari, a mummy from the same site (***19***). Bottom, the preserved leaf and the sequenced fragment seen from below (*inset*, magnification 250×). **(C)**, Papyrus de Kamara (***30***), illustrating a *Citrullus* fruit (*inset*), interpreted as a wild watermelon by Keimer (***20***). (**D-E**), Wall illustrations from Pharaonic tombs. (**D**), Tomb of Chnumhotep, Saqqara, *ca*. 4450 BC. (**E)**, Unspecified tomb from Meir, drawing by L. Manniche. Photo and drawing courtesy of L. Manniche, June 2018.

That the Egyptians ate domesticated watermelons is suggested by at least two wall paintings (**Fig. 1C-D**; *Supplementary text*) as well as *Citrullus* leaves from a tomb at Deir el-Bahari near Luxor (Theban Tomb number 320) first identified by the German archaeobotanist Georg August Schweinfurth in the 19^th^ century (*10, 11,19-21*) (**Fig. 1B**). The leaves were laying on the mummy of “Unknown Man C” in a re-used coffin inscribed for a priest Nebseni. On the evidence of embalming techniques and the coffin type the leaves may date to the early 18^th^ Dynasty (c. 3493-3470 BP; see *Supplementary text*). Finding leaves associated with mummies, however, is insufficient evidence that Egyptians consumed raw, sweet watermelons, since pharaonic tombs contain many kinds of plants that were not eaten as food, instead used as medicine or for other purposes (*10*). The closest known relative of the modern watermelon is a form from West Africa (*16, 17*) with white, usually bitter or bland pulp, and seeds that are used in West African ‘egusi’ stews (*18*).

To determine whether ancient Egyptians already consumed sweet, red watermelons and test whether sweet white-fleshed Sudanese *Citrullus* populations represent potential watermelon progenitors, we generated skimmed genome data (∼14 Gb) for all *Citrullus* species, including the Unknown Man C mummy sample (∼8 Gb) and accessions from North and South Darfur (Supporting Information **Table S1**) hypothesized to be the wild progenitor of watermelons (*6, 9-15*). The specific goals were to generate a strongly supported phylogeny, probe the phylogenetic placements of the Unknown Man C mummy sample and Kordofan melons, and searched bitterness and red-flesh markers in the ∼3500-year-old sample.

## Results and discussion

The 3500-year-old priest Unknown Man C mummy melon yielded good quality DNA with relatively low rates of deamination (SI Appendix **Figs. S1, S2**). Because aDNA yields short reads, we followed approaches used in other aDNA studies (*22, 23*) by mapping read data against target sequences from the watermelon genome (*16*). The targets consisted of plastid genes, a set of 143 nuclear genes for the phylogenomic analyses, and a set of 73 genes of interest linked to fruit bitterness and color. We recovered a complete plastid genome for the 3500-year-old leaf (SI Appendix **Fig. S3**), and its nuclear sequences were sufficiently good so that we could recover both our target genes for phylogenomics and our genes of interest related to cucurbitacin and lycopene pathways. This enabled to probe the phylogenetic position of the 3500-year-old *Citrullus* sample, and test whether its fruits were bitter or sweet, and red or white-fleshed.

For the phylogenetic analyses of *Citrullus*, we assembled entire plastid genomes (121 genes and 33 spacers) and 143 single copy nuclear genes (**SI Appendix, dataset 1**) and confirmed that nuclear genes are orthologs in reciprocal best alignment heuristic searches (*Materials and Methods*, **SI Appendix, dataset 2).** Maximum Likelihood (ML) nuclear and plastid phylogenies revealed the same topology (SI Appendix, **Fig. 2A; Fig. S4**), and a backbone similar to an earlier phylogeny based on 11 genes that lacked any East African samples (*17*). A coalescence-based tree also supported the same topology as ML analyses (SI Appendix **Fig. S5**). These phylogenomic analyses revealed that the Unknown Man C mummy sample is sister to modern domesticated watermelon (**Fig. 2A;** SI Appendix **Figs. S4, S5**) and both together are sister to the Sudanese sweet and white-fleshed Kordofan melons (**Fig. 2A**; *C. lanatus* subsp. *cordophanus*). All three form the sister clade to the West African *C. mucosospermus* (**Fig. 2A**). These analyses revealed that the 3500-year-old Unknown Man C mummy sample indeed represents a cultivated watermelon and that Kordofan melons (*C. lanatus* subsp. *cordophanus*) are the closest known relative of the watermelon and likely descendants of a progenitor population of the domesticated watermelon. All *Citrullus* species are illustrated in **Fig. 2B-I**.

**Fig. 2.**
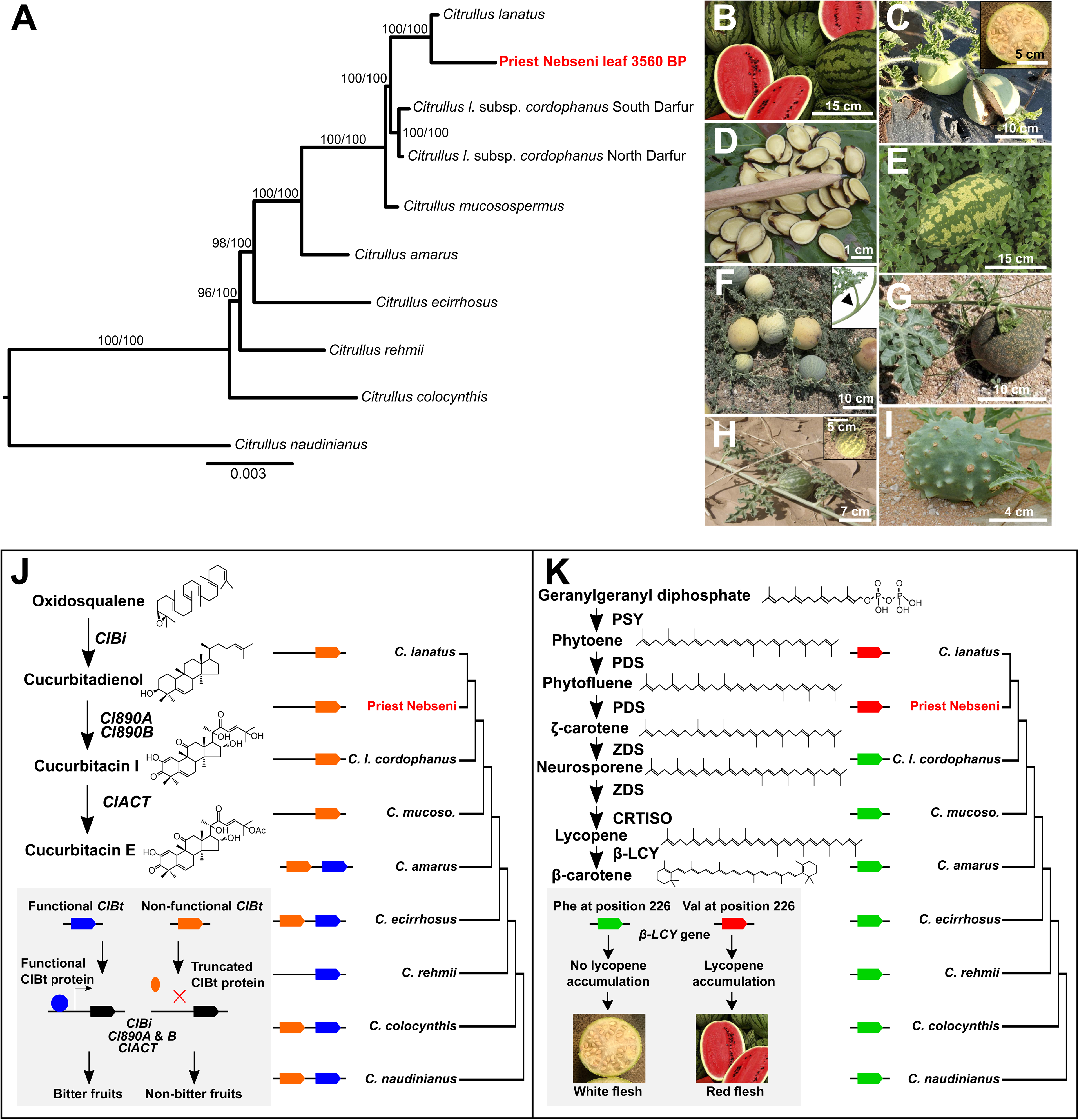
Phylogenomics of *Citrullus* reveals the watermelon progenitor and confirm that New Kingdom leaves belong to *Citrullus lanatus*. **(A)** Phylogeny of all *Citrullus* species based on 143 low copy nuclear genes, rooted on the relevant outgroups (***17***) (see also SI Appendix, **Figs. S1, S2**). Values on branches are bootstrap support from 1,000 replicates under the same ML model applied to nuclear (left) or plastid DNA sequences (right). **(B-I)**, Illustrations of all *Citrullus* species. **(B)** *Citrullus lanatus* (watermelon). **(C)** *C. lanatus* subsp. *cordophanus* (Kordofan melon). **(D)** *C. mucosospermus* (egusi melon). **(E)**, *C. amarus* (citron melon [wild], preserving melon [domesticated]). **(F)** *C. ecirrhosus* (tendril-less melon). **(G)**, *C. rehmii*. **(H)** *C. colocynthis* (colocynth). **(I)** *C. naudinianus*. (**J-K**) Key changes in genetic pathways underlying bitterness and lycopene synthesis in *Citrullus*. (**J**) Cucurbitacin pathway in *Citrullus* highlighting key changes in copy number and functionality of the fruit-specific transcription factor *ClBt* that controls fruit bitterness. (**K**) Lycopene pathway in *Citrullus*, and phylogenetic distribution of a key substation in LYCB (lycopene β-cyclase) gene that is linked to lycopene accumulation and controls fruit color.

To find direct evidence of the timing of watermelon domestication, we next investigated whether the 3500-year-old leaf belonged to a watermelon with bitter *vs*. sweet fruits and white *vs*. red flesh. Bitterness in cucurbits depends on terpene compounds called cucurbitacins. The *Bi* gene, which encodes an oxidosqualene cyclase (*OSC*) that catalyzes the first committed step in cucurbitacin C biosynthesis is critical for determining bitterness (*24-26*). Two bHLH transcription factors (*Bl* and *Bt*) regulate Cucurbitacin C biosynthesis by upregulating *Bi* expression in the leaves (*Bl*) and fruits (*Bt*) directly via binding to the E-box elements of the *Bi* promoter (*25, 26*). In addition to upregulating *Bi*, the watermelon *ClBl* and *ClBt* upregulate most other cucurbitacin metabolism genes, including the two cytochrome P450 enzymes that convert cucurbidienol in cucurbitacin precursor (*Cl890A* and *Cl890B* genes in watermelon) and the acyltransferase that convert the cucurbitacin precursor in cucurbitacin E (*26*). In cucumber (*Cucumis sativus*), honey melon (*Cucumis melo*), and watermelon, examination of different lines with varying bitterness revealed that domestication of non-bitter cucumber occurred via a nucleotide substitution leading to a premature stop codon in the *Bt* gene resulting in a truncated, non-functional protein (*24-26*). A genomic analysis comparing cucumber, honey melon, and watermelon (producing cucurbitacins C, B and E, respectively), revealed a domestication sweep at the *Bt* locus, with the loss of bitterness being due to convergent mutations at this locus (*26*).

We used a comparative-genomic approach to compare genes in the cucurbitacin biosynthetic pathway and its regulation across *Citrullus* species and the ancient watermelon. All cucurbitacin metabolic genes were conserved, with the same copy number across all *Citrullus* species, including the 3500-year-old sample (**Supplementary Dataset S1**). To probe whether the ∼3500-year-old watermelon had bitter or sweet fruits, we compared its *Bt* gene to that of the other *Citrullus* species. Most *Citrullus* species have two copies of *ClBt*, with the exception of *C. rehmii* and the clade comprising *C. mucosospermus*, the Kordofan melon, and the modern and ancient watermelon, which have a single copy (**Fig. 2J**). The second copy appears non-functional (**Fig. 2J, Supplementary Dataset S1**). The analyses further revealed that the 3500-year-old mummy sample has a single nucleotide substitution leading to a premature stop codon in the *Bt* gene (**Fig. 2J**, SI Appendix **Fig. S6A and dataset 1**), implying that it had non-bitter fruits. The Kordofan melons (subsp. *cordophanus*) also have the premature stop codon, matching their consistently sweet pulp (*13*). The fruits of the West African *C. mucosospermus* are usually bitter, but over 20% have acid, plain, or sweet pulp (*18: Table 5*), matching our finding of a stop coding in its *Bt* gene. The particular plant that we sequenced, *E.G. Achigan-Dako 809AA603* (SI Appendix, Table S1), indeed had non-bitter pulp (*18*; Achigan-Dako, pers. comm. SSR, 22 Nov. 2018). These results suggest a scenario wherein loss of pulp bitterness was a pre-adaptation for the domestication of watermelon, implying that early farmers probably took into cultivation non-bitter plants from the wild. Thus, our comparative analyses of *ClBt* genes across all *Citrullus* species suggests that the loss of bitterness resulted from initial selection of a non-bitter wild from, rather than selection occurring during cultivation as may have been the case in cucumber (*25, 26*).

The red pulp color in modern watermelon is due to lycopene accumulation, likely by blocking the conversion of lycopene into β-carotene, a step mediated by the enzyme LCYB (lycopene β-cyclase) (*16, 27, 28*). LCYB is encoded by the *lcyb* gene, and watermelon accessions with red flesh are characterized by having a unique A -> C substitution in the *lcyb* gene on chromosome 4, resulting in a Valine instead of a Phenylalanine at position 226 (*27*). To determine whether the 3500-old watermelon was red-fleshed, we sought all lycopene metabolic genes across our *Citrullus* genomes. This revealed a high conservation of these metabolic genes, with the same copy number for all genes in the lycopene pathway across all species (**Supplementary Dataset S1**). We found that the 3500-year-old mummy sample and the modern watermelon share the key substitution in LCYB linked to lycopene accumulation (**Fig. 2K, SI Appendix Fig. S6B and SI dataset 1**), while all other *Citrullus* species including the Kordofan watermelons lack this mutation and have white to greenish pulp. This reveals that New Kingdom Egyptians were cultivating a red, sweet watermelon 3500 years ago.

## Conclusion

Our study reveals that New Kingdom Egyptians cultivated a red, sweet watermelon at least 3500 years ago. It also shows that likely progenitor populations still exist in the upper Nile valley, in what is now the Darfur region of Sudan. The sweet, red watermelon may have been domesticated there and the use of this plant then could have spread northward along the Nile valley. Our finding that the 3500-year-old leaf belonged to a red-fleshed and sweet watermelon is consistent with two Egyptian wall paintings dating back to 4350 BP that have been interpreted as showing watermelons (**Fig. 1C-E**; *Supplementary text*). They show elongate fruits with the characteristic white stripes of watermelons and were served on trays, suggesting they were eaten raw. Watermelon may have been domesticated even earlier, given a watermelon seed found at Uan Muhuggiag in the Tadrart Acacus area in Libya that is between 5000 and 8650 years-old (*29*). Uan Muhuggiag lies 2240 km west of the 18^th^ Dynasty Theban necropolis of Deir el-Bahari where the genomic data here produced originated. Recent work on barley (*22*) and maize (*23*), and the present study on the watermelons, exemplifies the power of integrating collections-based phylogenomics with archaeological data and comparative genomics to resolve the domestication history of modern crops. The search for wild watermelon germplasm for engineering more disease-resistant varieties now should concentrate on the Darfur region of Sudan.

## Methods

### Plant taxon sampling

Material sequenced for this study is listed in **Supporting Information Table S1**, which also reports GenBank accession number for the skimmed genomes. *Citrullus* taxonomy follows Chomicki & Renner (*17*) and Renner et al. (*6*). We sampled all seven *Citrullus* species using herbarium-verified samples all of which are linked to permanently kept vouchers. This is essential, since very few –if any– germplasm accessions (for instance, *Citrullus* accessions in the National Plant Germplasm System Germplasm Resources Information Network [GRIN; http://www.ars-grin.gov/npgs]) are linked to herbarium voucher specimens, leading to problems of unverifiability (see for instance Renner et al. (ref. *6*)).

### Ancient DNA extraction

To document the leaf fragment received from Kew’s economic plant collection, we first took high resolution photographs (**Fig. 1B**). A small piece of a leaflet (<1 cm^2^, 5.8 mg) was ground with a Retsch mill (Retsch MM 400). DNA extraction was performed following Wales et al. (***31***) (modified by Pedersen et al. ref. *32*) and Dabney at al. (*33*). For the digestion treatment, a lysation buffer containing 0.5 % (w/v) N-Lauroylsarcosine (Sigma Aldrich L9150-50G), 50 mM Tris-HCl (Thermo Fisher Scientific 15568025), 20 mM EDTA (VWR E177-500MLDB) 150 mM NaCl (Thermo Fisher Scientific AM9760G), 3.3 % 2-Mercaptoethanol (Sigma Aldrich 63689-25ML-F), 50 mM DL-Dithiothreitol (Sigma Aldrich D9779-250MG) and 0.25 mg/mL Proteinase K (Promega V3021) was applied to the leaflet powder as described in Wales et al. (***31***). DNA purification was performed according to Dabney at al. (***33***) with reduced centrifugation speed (450 × *g*) as described in Basler et al. (***34***).

### Ancient DNA library preparation and sequencing

DNA extract was converted to an Illumina sequencing library using the single-stranded approach described in Korlević et al. (*35*). The protocol included the treatment with Uracil-DNA-Glycolase (New England Biolabs M0279) to remove Uracil residues and Endonuclease VIII (New England Biolabs M0299) to cleave DNA strands at abasic sites. 2.5 U/µl of Circligase II (Biozym 131406) was used for the fill-in reaction and carried out overnight. A quantitative PCR was performed on a PikoReal 96 Real-Time PCR machine (Thermo Fisher Scientific TCR0096) using 0.2 % of the unamplified library following this thermal profile: 10 min initial denaturation step at 95 °C, followed by 40 cycles of: 15 s at 95 °C, 30 s at 60 °C, and 1 min at 72 °C. The quantitative PCR reaction mix contained a final volume of 10 µL: 1 µL of diluted library, 1 × SYBR Green qPCR Master Mix (Applied Biosystems 4309155), 0.5 µM of each primer IS7 and IS8. Three replicates of each library were used. Indexing PCR was performed by the appropriate number of cycles adding 8 bp indices to 5’ and 3’ adapters. PCR and final concentrations are the same as described (*36*), but with a final volume of 80 µL using 20 µL of template. DNA sequencing was performed on an Illumina NextSeq 500 sequencing platform, using the 500/550 High Output v2 kit (75 cycles, Illumina FC-404-2005), with a custom read-1 (*36*) and a custom index-2 (*37*) sequencing primer. Extraction and library building was performed in the ancient DNA facility of the University in Potsdam; negative controls were included in all steps.

### Genome skimming of *Citrullus*

We extracted genomic DNA from all *Citrullus* samples except the 3500-year-old sample using the Quiagen DNeasy Plant kit, following the manufacturer’s protocol. Genome skimming was performed by the company Genewiz (New Jersay, USA) in Germany. Paired (2×150 bp) DNA libraries were sequenced on an Illumina HiSeq platform and 350 GB of sequence data were produced. An average of 100 million reads were produced for each sample.

### Plastid data processing, assembly and annotation

The Illumina raw reads were quality filtered using Trim Galore v.0.4 (*38*), discarding sequences with averaged phred33 score below 20. Pre- and post-trimming read quality was assessed using FASTQC v.0.1 (*39*). Whole plastid genomes were assembled by blasting quality-filtered reads against a fully annotated reference plastid genome of *C. lanatus* (GenBank accession NC032008) using the tool *blastn* and a maximum number of target sequences of one. We obtained the best scoring blast hits from the blastn output by filtering all hits by bitscore and query coverage values of 100 and 80%, respectively. Best scoring reads were mapped against the reference plastid genome using the Geneious mapping tool (*40*) and the following parameters: five iterations, minimum mapping quality of 30, maximum gap size of 100, and maximum mismatches per read of 40%. Consensus genome sequences were assembled following a modified statistical base-calling approach of Li et al. (*41*), i.e. minimum depth coverage of 10, and bases matching at least 50% of the reference sequence. Plastid genomes were fully annotated by transferring intron, exon, and spacer annotation from the reference plastid genome to the *de-novo* assembled plastids.

### Plastid and nuclear phylogenomics of *Citrullus*

De-novo plastid genomes were aligned with Mauve using a progressive algorithm and assuming collinearity (*42*). The resulting ∼150,000 bp alignment was subjected to Maximum Likelihood (ML) tree inference in RAxML v8.0 (*43*), using the GTR substitution model, 25 gamma categories, and 1,000 bootstrap replicates. Nuclear phylogenomic inferences relied on a selection of 143 out of 353 nuclear CDS known to be present in low copy numbers across angiosperms (*44*), all assembled from the trimmed read data using the python package Hybpiper (*45*). The 143 genes (**SI dataset 1**) were selected based on the proportions of missing data (i.e. alignments including more than five sequences with 50% of their length missing and less than five taxa were excluded), and phylogenetic informativeness (PI) profiles (i.e. gene alignments with disproportionate substitution rates were excluded) following Townsend (*46*). The PI profiles are provided in SI Appendix, **Fig. S7**. Hybpiper-achieved gene assembly by mapping the trimmed reads against the set of targeted CDSs using the Burrows-Wheeler alignment software (*47*) with a coverage threshold of 8x. Contig assembly relied on the software SPAdes (*48*), relying on default settings. Whenever multiple copies of the target genes are present in the genome, HybPiper raises paralog warnings during the contig assembly step. Here, no paralog warnings were issued for any of the 143 low copy nuclear genes. Because the Illumina sequencing of the ancient *C. lanatus* sample produced very short reads (max length of 75 bp; SI Appendix, **Fig. S1**), we carried out low-copy-gene assembly using the Geneious mapping algorithm, using customized settings. Gene mappings and assemblies were all checked by eye and carefully curated. To determine whether the selected set of low copy nuclear genes are in fact orthologs, we carry out orthology analyses in the software ProteinOrtho (*49*), which relies on a reciprocal best alignment heuristic approach. No paralogues were detected by ProteinOrtho (**SI Appendix, dataset 2**).

Phylogenomic analyses of the unlinked gene alignments matrix relied on ML and coalescence-based methods. We generated ML trees for every nuclear gene alignment as well as the concatenated nuclear matrix using the same settings as employed for plastid tree inference. Species tree analyses relied on the coalescence-based method ASTRAL III (*50*), using as input the annotated consensus gene trees derived from RAxML, with branches having Likelihood Bootstrap Support (LBS) < 10 collapsed. Gene tree conflict was visualized by plotting the local posterior probabilities derived from ASTRAL as pie diagrams at nodes. The final normalized quartet score is: 0.73.

### In-silico capture of genes relating to cucurbitacin biosynthesis and its regulation, lycopene synthesis and sugar metabolism and its regulation in *Citrullus* species

We followed the same read-data filtering and trimming procedure as used for the plastid data. Nucleotide deamination analysis and filtering of the ancient watermelon reads were carried out by the pipeline PALEOMIX (*51*) and using as a reference a genome assembly of a modern accession of *C. lanatus* (No. 97103, available at http://cucurbitgenomics.org/). The pipeline relied on the software Bowtie2 (*52*) for read mapping and on mapDamage2 (*53*) for nucleotide deamination analysis. Genes involved in the cucurbitacin and lycopene synthesis were also captured in-silico following the same approach employed to data-mine low copy nuclear genes in the pipeline HybPiper. To attain this, we build a customized set of target genes obtained from Guo et al. (*16*) and Zhou et al. (*26*) (gene IDs are provided in **SI Appendix, dataset 3**). Gene copies in the *ClBt* gene were detected by HybPiper during the contig assembly step. No paralog warnings were produced for the remaining genes involved in the cucurbitacin and lycopene pathways (except for the *ClBt* gene for which some species had two copies– cf. *Results and discussion*).

## Acknowledgements

We thank the egyptologists, R. Schiestl, University of Munich, Richard Parkinson, University of Oxford, and John H. Taylor, Department of Ancient Egypt and Sudan, The British Museum, for consultation about the age of the mummy; and S. Bellot (Royal Botanic Gardens, Kew) for advice on *in-silico* gene capture. This project was supported by the German Science foundation (DFG grant RE 603/27-1) and the Elfriede and Franz Jakob Foundation for research in systematic botany. G.C. is supported by a Glasstone Fellowship and a Junior Research Fellowship at the University of Oxford. O.A.P.E. is supported by the Lady Sainsbury Orchid Fellowship.

## Author Contribution

SSR and GC conceived the study with subsequent methodological contribution from OAPE; SSR, MVS, MP gathered data; OAPE, GC, SSR analyzed data, SSR, GC, MH provided funding; SSR and GC wrote the manuscript with edits from all authors.

## Additional information

**Supplementary Information** accompanies this paper at https: xxx

## Figure legends

**SI Appendix consists of SI text, Table S1, SI figures 1-7, and supplementary datasets 1, 2, and 3**. All DNA sequence alignments generated for the phylogenomic analyses as well as for the genes of interests. Dryad accession number forthcoming.

