## Supplementary material for "A 3500-year-old leaf from a Pharaonic tomb reveals that New Kingdom Egyptians were cultivating domesticated watermelon": SI Appendix

**Supplementary text**

***Three main hypotheses on the origin of the watermelon***

**The hypothesis of a South African origin** of watermelon originated only in the 1930s, when Bailey (1) erroneously synonymized the South African species *C. lanatus*, collected near Cape Town in 1773 and described by Thunberg, with Linnaeus’s domesticated watermelon, which is based on a cultivated specimen collected in Southern Italy. DNA sequences of Thunberg’s specimen as well as a dense sampling of all other species of *Citrullus* have demonstrated that the domesticated watermelon and the material from South Africa are not closely related (2). Instead, the South African type material is sister to accessions of *C. ecirrhosus*, an endemic species of the Namib-Kalahari. *Citrullus* now contains seven species, of which four are endemic in South Africa: *C. amarus* (a replacement name for *C. lanatus* (3)), *C. ecirrhosus*, *C. naudinianus*, and *C. rehmii* from the Namib-Kalahari region, *C. colocynthis* from northern Africa to west India (now naturalized in Australia) and *C. mucosospermus* from Benin, Ghana, and Nigeria. The rejection of a South African origin of the watermelon matches lack of any archeobotanic evidence and the absence of early farmers there (4).

**The hypothesis of a West African origin** goes back to molecular phylogenies that showed watermelon as sister to the West African *C. mucosospermus* (3, 5). However, *C. mucosospermus* is not well represented in germplasm collections, which has led to the inclusion of misidentified accessions supposed to represent the species. Thus, of the six accessions of this species re-sequenced by Guo et al. (5) only PI249010 from Nigeria may represent *C.* *mucosospermus*, while PI482271 from Zimbabwe is *C. amarus* (*6*), PI189317 from Nigeria is *C. lanatus* (7), PI500301 from Zambia and PI248178 from Zaire are from outside the known range of the species (8), and so is PI595203 from Georgia, USA. Sequencing of herbarium-verified *C. mucosospermus* accessions nevertheless supported Guo et al.’s finding of a sister relationship between watermelon and *C. mucosospermus* (2). Both these studies, however, failed to include potential wild populations of watermelon from East Africa. There is also no archeobotanic evidence of early melon farming from West Africa.

**The hypothesis of an East African, specifically Upper Egyptian, origin** of the watermelon originated with Schweinfurth (9-12) who was the first botanist to collect watermelons throughout Egypt and Sudan and to identify plant material discovered in the Pharaonic tombs when asked by the director of the Cairo museum, G. Maspero (1846-1916) to sort and distribute botanical material from the tombs to European collections (12). He was consulted during the unpacking of the mummies of Ramesses II, Ahmose I, and others from the 18^th^ or 20^th^ dynasties found in the Theban Necropolis near Luxor. In 1869, Schweinfurth discovered watermelons with globose, small, but edible fruits south of Khartoum on the White Nile (‘Kordofan’ or Kurdufan today Darfur, Sudan) that he identified as a wild form of *C. vulgaris* (the accepted name of the domesticated watermelon before Bailey’s (1) wrong synonymization), and in 1882, when collecting between Asyut and Aswan, he noted melons with inedible and even smaller fruits (10-14 cm in diameter) that he called *C. vulgaris* var. “*colocynthoides*”, in reference to the perfectly globose, small, and inedible fruits of *C. colocynthis*. Both these Schweinfurth collections, consisting only of seeds, are now in the Berlin herbarium, and nuclear and plastid DNA sequences obtained from them indicate that the Khartoum plant represents a form of watermelon while the Asyut plant (with 10-14 cm fruits) is *C.* colocynthis (13). This matches the colour, shape, and testa surface of the seeds: colocynth seeds are more convex, pointed, and smoother, while watermelon seeds are slightly larger, have a rougher surface, and have tiny wings at the base. The common Arabic names for the colocynth are gurma or gurum, but the same names apply also to the small-fruited, edible wild watermelons from the Kordofan region (11, 12). While watermelon is an annual, the colocynth is a perennial plant; it is still cultivated today for its seeds, which are eaten toasted (often salted) or are used for oil extraction, while the fruit pulp, which is white and bitter, is discarded (14).

***Oldest records for Citrullus species and dating of the leaf sequenced in this study***

**The oldest records of *C. colocynthis*** come from the Neolithic site Uan Muhuggiag in southwest Libya, in layers radiocarbon dated to 8620 and 8650 BP; the seed itself was not dated, and stratigraphic evidence suggests that the layer was disturbed and contains younger intrusive material (15). The same layer also included one seed of *C. lanatus* (15). Another single *C. colocynthis* seed was found at Helwan near Cairo, a site dating to 6000 BP (16), and several seed fragments in a tomb from Ne-user-re (Niuserre, Neuserre) in the necropolis of Abusir dating to 4455-4420 (17).The gut of a predynastic mummy from Naga-ed-Der dated to the Middle Kingdom, 7300–3650 BP, contains seeds of *C. colocynthis*, as drawn and identified by Schweinfurth and reported by Netolitsky (18) (the drawing is reproduced as Fig. 1 by Wasylikowa and van der Veen (15)), and so did the stomach of the mummy of Ramesses II, dated to 2300 to 3200 BP [died c. 1212 BC] (Germer (19), p. 9).

**The oldest record of *C. lanatus*** is the single seed from Libya (see *main text* but whether this seed came from a domesticated watermelon is unknown), and younger seeds have been reported from an 18^th^ dynasty temple (3500-3480 BP) near Semna in Sudan (20), but without morphological details or illustrations, and the material has been lost (email from Sylvia Blomsma, Research Assistant at Palaeobotany Dept., Groningen Institute of Archaeology, University of Groningen. 30 March 2017). Well-preserved *C. lanatus* seeds were found by H. Carter and P.E. Newberry in two oval baskets in Tutankhamun’s tomb at Thebes, dated to around 3350 BP. Some of these are kept at Kew (accession H. Carter 4.1965), some in the Dokki Agricultural Museum in Kairo (accession 4198; photograph in Darby et al. (21), Fig. 18.8, reproduced as Fig. 1.4 in Gyulai et al. (22)). “They were still in remarkably good condition as flat seeds about 8 mm (5/16 in) long. When fresh they would have been lightly roasted and then eaten by cracking the brittle outer cover with the front teeth” (Hepper (23) p. 56, plate 34; also Andrews (24)).

The **only known ancient leaves of *Citrullus*** come from Theban Tomb number 320 at Deir-el-Bahari near Luxor discovered in 1881, where they were laying on the mummy of an unknown man, referred to as the priest Nibsoni or Nebsoni (10, 11, 25-27: mummy CG61067, Unknown Man C). Precise dating is problematic because the 20^th^ to 18^th^ dynasty tombs of Deir-el-Bahari were repeatedly disturbed by grave robbers who mixed up mummies and coffins; conservatively, these tombs date to the 18^th^ dynasty, 3550-3290 BP (27a = Ikram and Dodson, 1998; R. Schiestl, Egyptologist, University of Munich, July 2017). Based on the evidence of the embalming techniques, the mummy of Unknown Man C is generally thought to date to the 18^th^ Dynasty (Ikram and Dodson, 1998: tentatively suggest c. 1450 BC). Other floral remains found in TT 320 have been radiocarbon dated: A lotus flower of Ramesses II (d. 1212 BC) has been dated to 1000-840 BC, a willow wreath from Ahmose I (d. 1515 BC) to 1381-1091 BC, plant material from Pinedjem II (d. 982 BC) to 1045-858 BC, in this case a match between the death of the pharaoh and the radiocarbon date of the plant material (27b = McAleely, 2013). The tomb was finally sealed c. 930 BC, and coffin number 320 was immediately inside the entrance, thus one of the last ones to be placed inside. The similar dating of the coffin and the mummy increases our confidence in the 18^th^ Dynasty date for the watermelon leaves sequenced in this study. The latest possible date for the *C. lanatus* leaf material is c. 930 BC or 2930 years before present.

Schweinfurth placed the leaves in water to flatten them and then compared them to both colocynths and the wild watermelons of Upper Egypt (above). “The problem to solve was whether the leaves were those of the water-melon or those of the colocynth, a species spread over the whole desert region and only differing from the former, which has long hairs on the young fruit, by the complete nudity and spongy nature of its bitter fruit with a hard rind and by the seeds. The leaves of the water-melon often very closely resemble those of the colocynth, especially in the variety called Gjurma (Gyurma) in Egypt, which bears fruit no larger than that of the colocynth, though it is always sweet.” (Schweinfurth (11), p. 113). After comparing long series of leaves of both species to the Pharaonic leaf, he continues “The leaves found on Nibsoni are about a palm long, and of a pinnatisect form, with obtuse leaves. If these leaves were distinctly hairy there would be no doubt of their belonging to the water-melon. Yet, as already mentioned, there is a variety widely spread in Egypt which has not the long and numerous hairs attached to the tubercles with which the leaves are covered, but merely short bristles, which is also the case in the colocynth. This variety of melon, which I have named *colocynthoides*, is the Gyurma of the Egyptians, and is cultivated in dry neglected ground in Upper Egypt. […] The leaves of the Gyurma are sometimes hairy as in the water-melon, sometimes only provided with short deciduous bristles, as in the colocynth.” Schweinfurth then sent the leaves to Hooker at Kew, where the accession book of 1883 lists “8 sheets of mounted specimens of mummy wreaths of flowers etc. from the coffins of Ramesses I and Ramesses II”, and a note with the leaf states “remains of leaves of *Citrullus vulgaris* Schrad. from the coffin of Nibsoni, a priest of the 20^th^ dynasty, No. 7, Dr. Schweinfurth.” (<http://apps.kew.org/ecbot/specimen/40730>).

In his numerous comments on watermelons spread over several papers, Schweinfurth also mentions the citron melon grown for making jams (its fruits are inedible raw and need to be cooked with sugar) saying, “La pastèque existe a l’état sauvage au Kourdofan, sur les bords du Nil blanc et même au pays de Niam-Niams où elle est a demi cultivée. Son fruit n’est pas plus gros que celui de la coloquinte.” (Schweinfurth (10), p. 201). At the time, the citron melon was considered a form of watermelon, but it is now known to be a separate species, *C. amarus* (2, 13). Still later (Schweinfurth (28), p. 73), he referred to the Nibsoni mummy leaves as belonging to the ‘pastèque’, “La pastèque est native de l'Afrique centrale où elle se trouve à l'état sauvage; le fruit en est moins grand et moins savoureux que celui de nos cultures, mais offre les mêmes caractères,” while in a third paper (Schweinfurth (12), p. 361), he returns to assigning the Nibsoni leaves to the wild gyurma watermelon of Aswan and Upper Egypt, *C. vulgaris* var. *colocynthoides.*

***Ancient Egyptian illustration of watermelon and colocynth***

So far, **two Ancient Egyptian illustrations of watermelon and one of a colocynth** are known. The first (reproduced in **our Fig. 1E**) shows a large oblong watermelon with black and white longitudinal stripes served on a tray and comes from a tomb from Meir, Northwest of Asyut (Manniche (29), p. 92; Janick et al. (30), Fig. 2A). The painting’s precise origin is not mentioned by Manniche (it is not from Blackman, 1914-1953; Manniche pers. comm. to S.S.R., 28 Feb 2018), but it probably dates to 4350-4200 BP (R. Schiestl, Egyptologist, University of Munich, February 2018). The second (**our Fig. 1D**) comes from the tomb of Chnumhotep near Saqqara, dated to 4360-4350 BP; 30a = Moussa and Altenmüller, 1977: pl. 89). Speculatively, the flat trays on which these fruits were served suggest that they were eaten raw, as a cold dessert, which would imply loss of the bitterness genes; however, being shown on a tray is not a definitive argument because, for instance animal horns were also sometime illustrated on a tray (R. Parkinson, pers. comm. to G.C. January 2018). The interpretation of these fruits as a desert fits the Ancient Egyptians’ popular saying “Fill up your stomach with a summer watermelon”, apparently meaning “Don’t worry” (Darby et al. (21), p. 217).

The illustration of the **colocynth** shows a perfectly globose fruit with black and white longitudinal stripes on a stalk with two leaves, slightly longer than the fruit. The drawing comes from a papyrus of the 21^st^ dynasty (31) and is reproduced by Keimer (26) (p. 170), who considers it the gyurma (wild) form of *C. vulgaris*. In its diameter relative to leaf length, perfectly globose shape, and conspicuous stripes is exactly matches photos of the wild watermelon, *C. lanatus* var. *cordophanus*, in Ter-Avanesyan (32).

Schweinfurth’s suggestion that he had found wild progenitors of watermelons caused the Russian breeder David Ter-Avanesyan (1909–1979) in the early 1906s to obtain seeds from the Kordofan region that were then planted in Tashkent at a research station of the Vavilov Center Sudanese (32) (Larisa Bagmet, Curator of VIR, pers. comm., 30 Mar 2017). These relatively small watermelons have non-bitter fruits with white pulp and a super aromatic taste, and both Ter-Avanesyan (32) and Fursa (33) considered this material to represent the progenitors of cultivated watermelon, therefore naming it formally as subspecies or varietas *cordophanus* of *C. lanatus*. An immunochemical analysis of seed proteins that included all species except *C. rehmii* (not yet discovered) bears this out (34).

27a. Ikram, S. and Dodson, A., 1998. The mummy in ancient Egypt: Equipping the dead

for eternity. London: Thames & Hudson, p. 316.

27b. McAleely, S. 2013. Garlands from the Deir el-Bahri cache. Pp. 153-166 in A.J.

Shortland & C.B. Ramsey, Radiocarbon and the chronologies of Ancient Egypt. Oxford: Oxbow.

28. Schweinfurth G (1882) De la flore pharaonique. *Bull Inst Egypt Ser 2* 3:51-76.
29. Manniche L & London British Museum. (1989) *An ancient Egyptian herbal* (pp. 133-134). Austin: University of Texas Press.
30. Janick J, Paris HS, Parrish DC (2007) The cucurbits of Mediterranean antiquity: identification of taxa from ancient images and descriptions. *Ann Bot* 100:1441-1457.

30a. Moussa, A. M. and H. Altenmüller. 1977. Das Grab des Nianchchnum und

Chnumhotep. Archäologische Veröffentlichungen 21. Mainz am Rhein.

31. Naville E (1912) Papyrus funéraires de la XXI dynastie. Vol. 1. Le papyrus hiéroglyphique de Kamara, le papyrus hiératique de Neskhonsou au Musée de Caire. Paris, Ernest Leroux.
32. Ter-Avanesyan DV (1966) Arbuz Kordofanskyi *Citrullus lanatus* Mansf. ssp. *Cordophanus* Ter-Avan. *Bot. Zhurnal* 51:423–426.
33. Fursa TB (1972) K sistematike roda Citrullus Schrad. [On the taxonomy of genus *Citrullus*] Schrad. Bot. Zhurn. (Moscow & Leningrad) 57:31–41.
34. Fursa TB, Gavrilyuk IP (1990) Phylogenetic relations of the genus *Citrullus* Schrad. based on the immunochemical analysis of seed proteins. *Sborn Nauchn Trudov Prikl Bot Genet Selekts* 133:19–26. [In Russian with English summary]

**Table S1**. Material used in this study. Nomenclature follows Renner et al. (2017). ‘(M)’ means that the respective herbarium voucher has been deposited in the Munich herbarium. Genome skimming datasets have been submitted to the Sequence Read Archive (SRA) of NCBI (<https://www.ncbi.nlm.nih.gov/sra>)

| *Citrullus* taxa | Source/voucher | Accession number |
| --- | --- | --- |
| *C. amarus* Schrader | S.S. Renner & M. Silber 2866 (M), grown in Munich from commercially bought seeds | Forthcoming |
| *C. colocynthis* (L.) Schrad. | A. Patzelt 4872 (M), grown in Munich from seeds collected in Oman | Forthcoming |
| *C. ecirrhosus*  Cogn. | S.S. Renner 2855 (M), grown in Munich from USDA Grif 16056 seeds originally from Namibia | Forthcoming |
| *C. lanatus* subsp. *cordophanus*  Ter-Avan. | M. Sir El Khatim 1 (M), grown in Munich from seeds originally from North Darfur, El Fashir (Al Fasher) | Forthcoming |
| *C. lanatus* subsp. *cordophanus* | S.S. Renner 2865 (M), grown in Munich from seeds originally from South Darfur, Adiela | Forthcoming |
| *C. lanatus* subsp. *vulgaris* (Schrad.) Fursa | M. Silber 19 (M), grown in Munich from commercially bought seeds of cultivar ‘Sugar Baby’ | Forthcoming |
| *C. lanatus* subsp. *vulgaris* (Schrad.) Fursa | Fragment of leaf from the coffin of Nebseni (‘Nibsoni’), leg. Dr. Schweinfurth, Kew Economic Plant collection  <http://apps.kew.org/ecbot/specimen/40730> | Forthcoming |
| *C. mucosospermus* (Fursa) Fursa | S.S. Renner & M. Silber 2869 (M), grown in Munich from Gatersleben seeds originally collected in Benin (E.G. Achigan-Dako 809AA603) | Forthcoming |
| *C. naudinianus* (Sond.) Hook. | J-L. Gatard s.n. (M), grown in Munich from commercially bought seeds, originally from Namibia | Forthcoming |
| *Citrullus rehmii* De Winter | S.S. Renner & M. Silber 2864 (M), grown in Munich rom commercially bought seeds, originally from Namibia | Forthcoming |
