## Supplementary material for "A 3500-year-old leaf from a Pharaonic tomb reveals that New Kingdom Egyptians were cultivating domesticated watermelon": Fig. S1

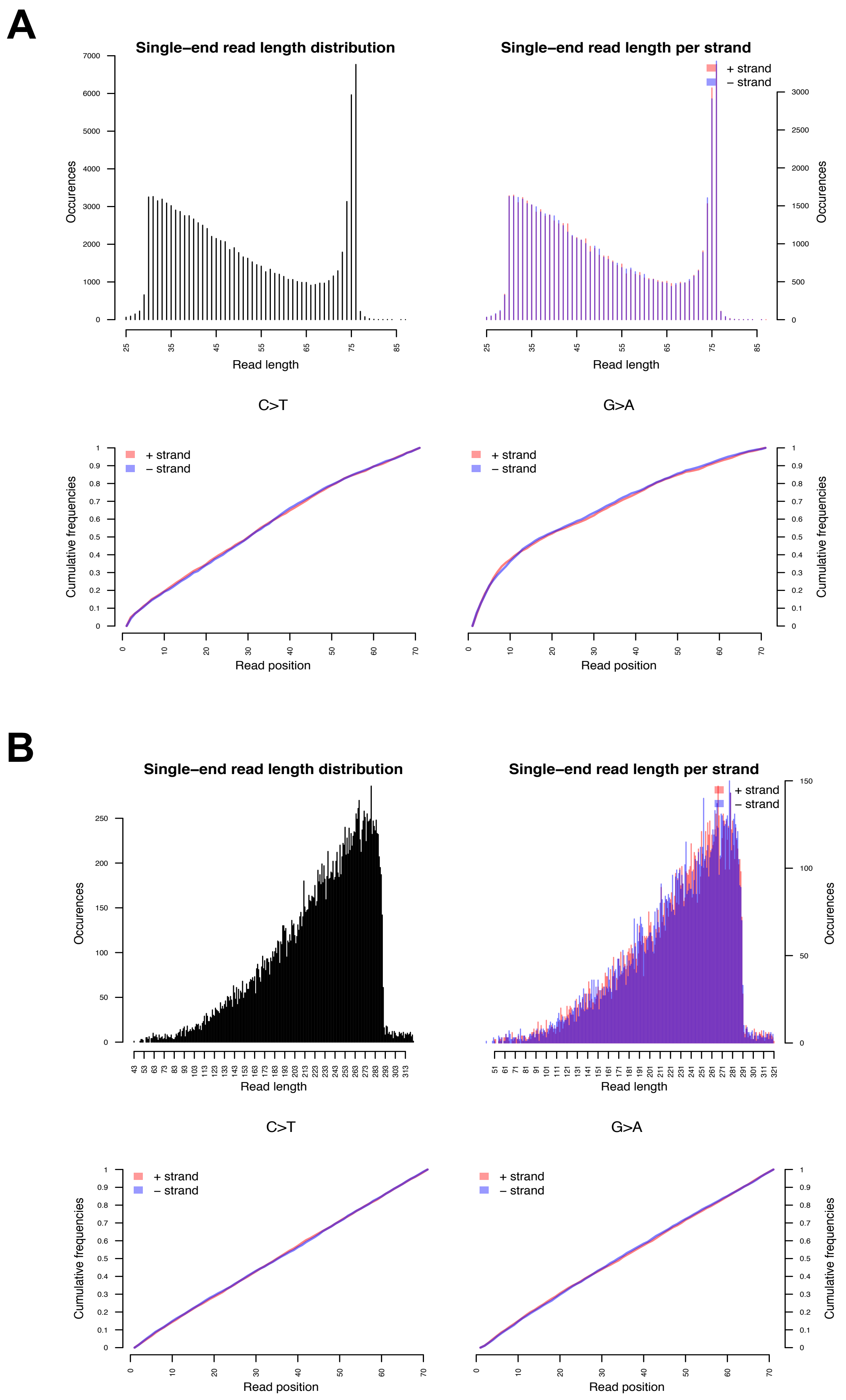


**Fig. S1.** Ancient *Citrullus* DNA sequencing. Distribution of read length in the ~3560-year-old *Citrullus* (**A**) compared to those of modern *Citrullus lanatus* (**B**). Cumulative frequencies of C > T and G > A changes in function of read position.
