## Supplementary material for "A 3500-year-old leaf from a Pharaonic tomb reveals that New Kingdom Egyptians were cultivating domesticated watermelon": Fig. S2

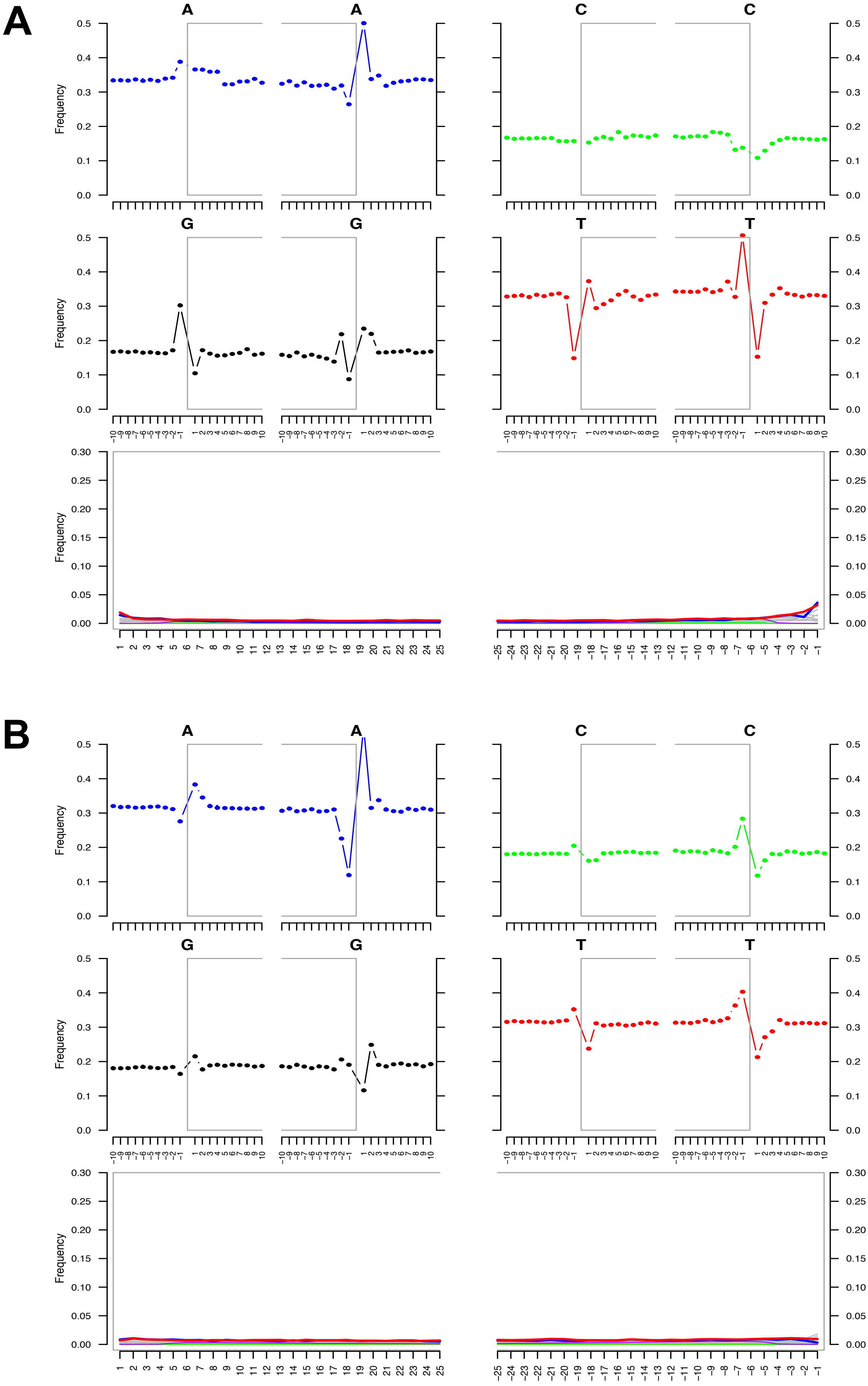


**Fig. S2** Authentification of *Citrullus* ancient DNA. Deamination patterns for each nucleotide in the Ancient *Citrullus* sample (**A**) compared to the modern accession of *Citrullus lanatus* (**B**). X axes indicate individual nucleotide positions of DNA fragments. In the four first plots in (**A**) and (**B**) 5' end is on the left and 3' is on the right.
