## Supplementary material for "A 3500-year-old leaf from a Pharaonic tomb reveals that New Kingdom Egyptians were cultivating domesticated watermelon": Fig. S3

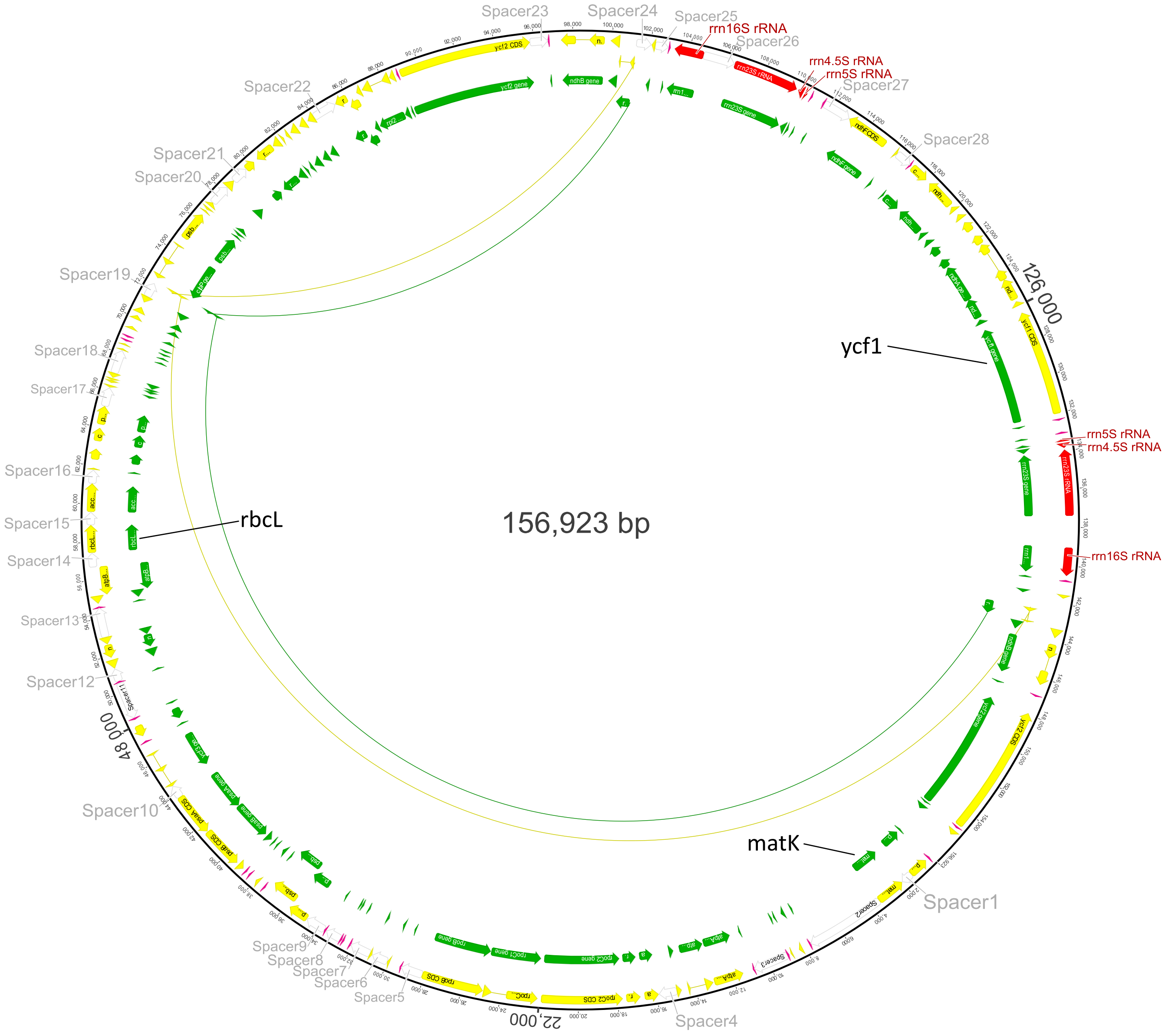


**Fig. S3.** The pastid genome of the ~3560-years-old *Citrullus* leaf is almost identical to that of modern *Citrullus lanatus*. The 158,923 bp plastome of the Pharaoh melon (green) was mapped against the 156,923 bp plastome of modern *Citrullus lanatus* (yellow).
