## Supplementary material for "A 3500-year-old leaf from a Pharaonic tomb reveals that New Kingdom Egyptians were cultivating domesticated watermelon": Fig. S4

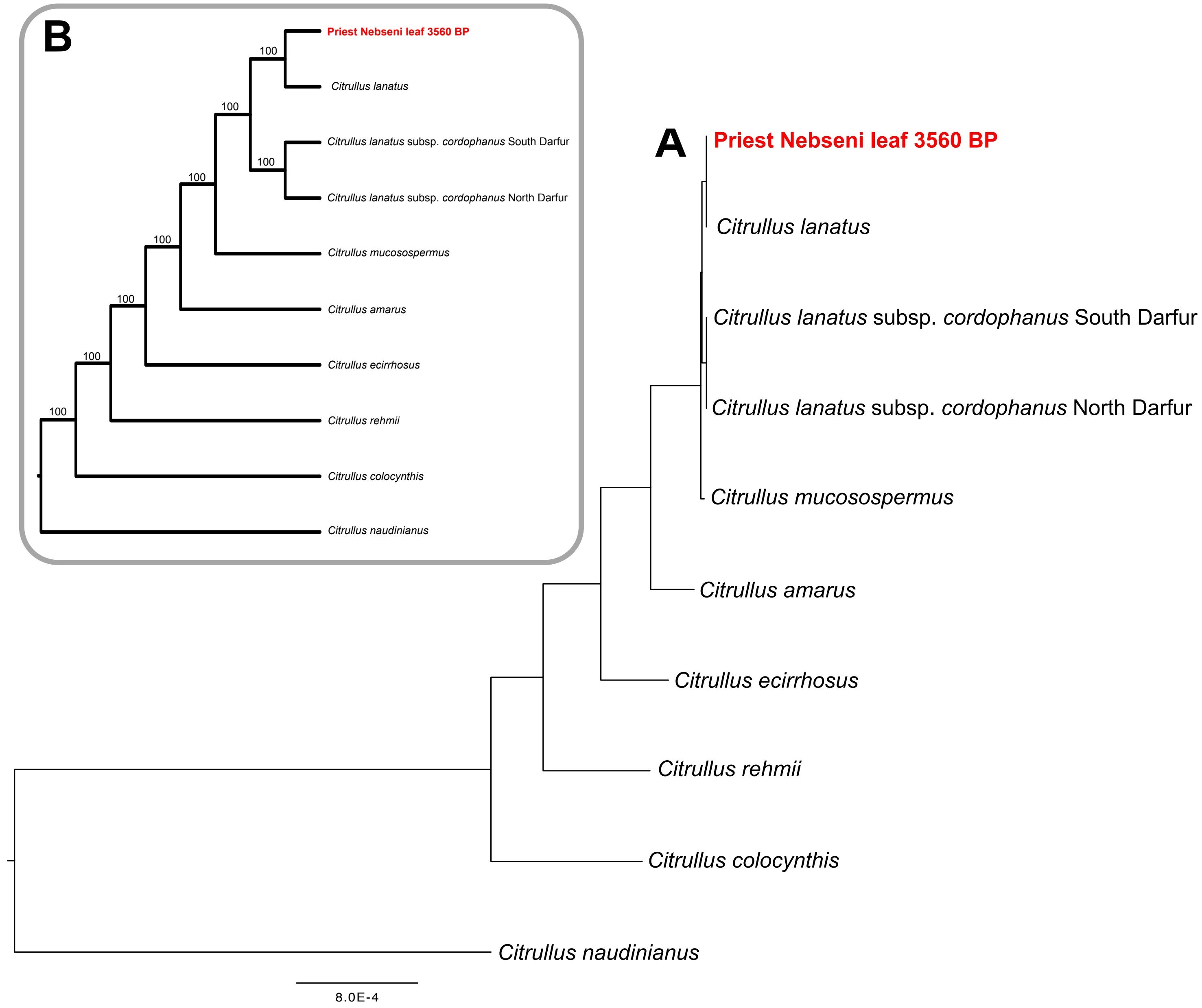


**Fig. S4.** Maximum likelihood phylogeny under a GTR + G model from *Citrullus* plastid genomes (121 genes and 33 spacers). (**A**) Maximum likelihood tree. (**B**) Cladogram showing the topology and bootstrap support from 1000 replicates under the same model as in (**A**).
