## Supplementary material for "A 3500-year-old leaf from a Pharaonic tomb reveals that New Kingdom Egyptians were cultivating domesticated watermelon": Fig. S5

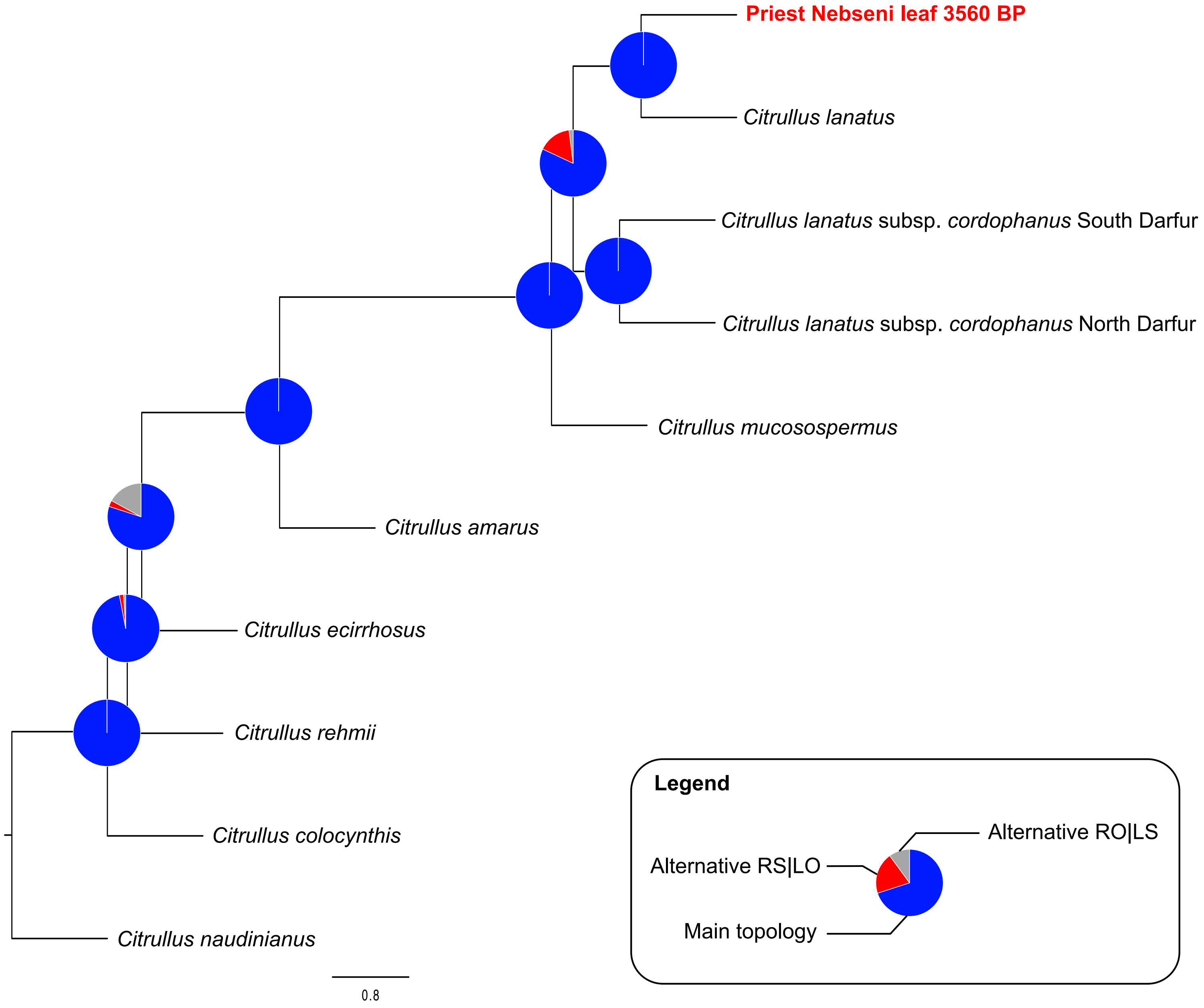


**Fig. S5.** Coalescence-based tree (Astral) for *Citrullus*, based on the 143 nuclear genes. The results show very little conflict between gene trees.
