## Supplementary material for "A 3500-year-old leaf from a Pharaonic tomb reveals that New Kingdom Egyptians were cultivating domesticated watermelon": Fig. S7

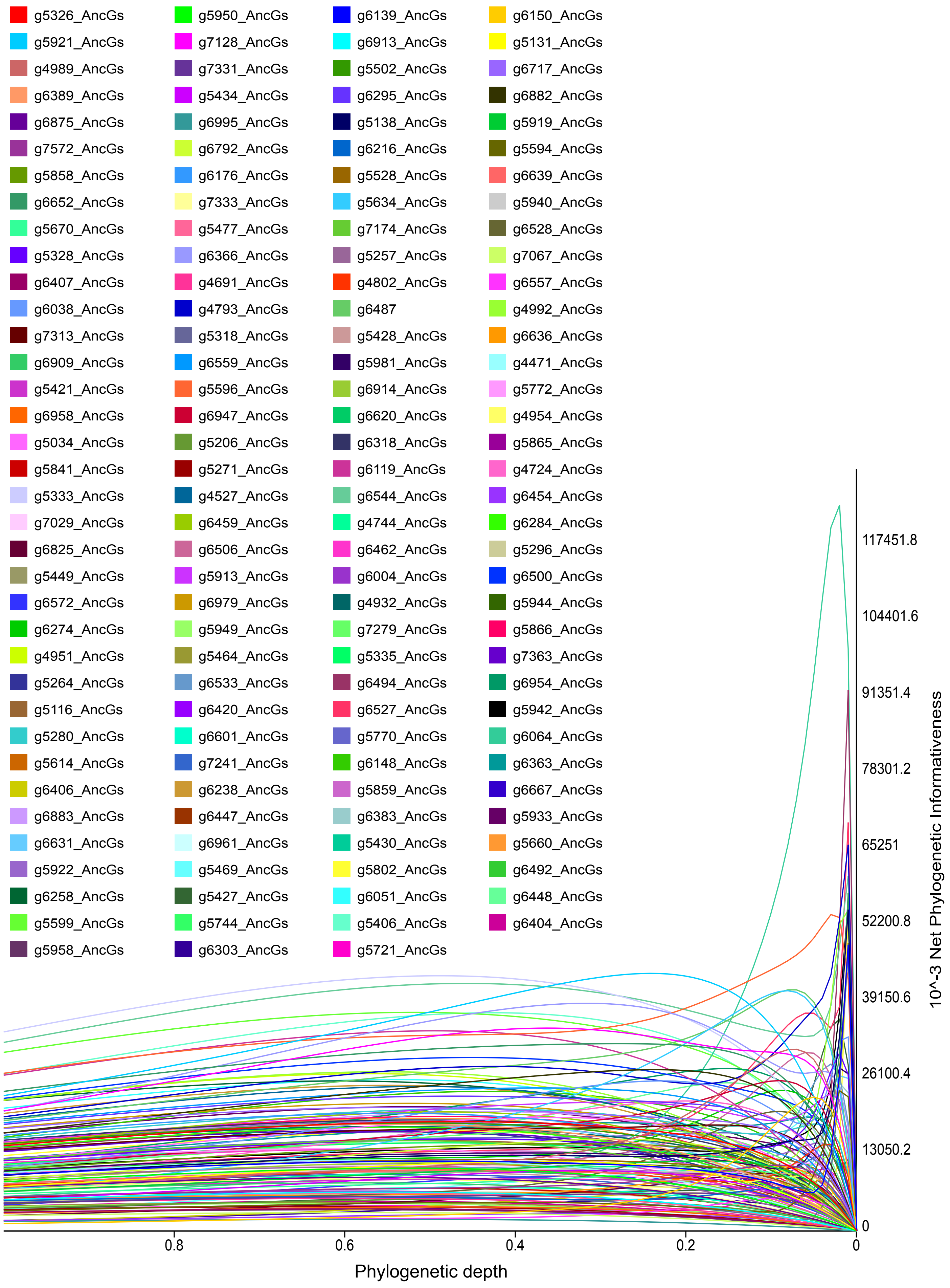


**Fig. S7.** Phylogenetic informativeness of the 343 nuclear genes (Johnson et al. 2018), based on which we selected the 143 nuclear genes used in our phylogenomic analyses. The gene color-code shown above reports the numbering system used by Johnson et al. (2018). For GenBank accession numbers of the gene targets, see Table S2.
